# Preferential recruitment of PML isoform II by incoming HPV16 genome transport vesicles facilitates early stage of infection

**DOI:** 10.64898/2026.11.09.765526

**Authors:** Anand Kushwaha, Malgorzata Bienkowska-Haba, Timothy R. Keiffer, Mireya Represa Perez, Rona S. Scott, Andrew D. Yurochko, Martin J. Sapp, Katarzyna Zwolinska

**Affiliations:** Department of Microbiology and Immunology, Center for Molecular and Tumor Virology, Center for Applied Immunology and Pathological Processes, Louisiana State University Health Sciences Center, Shreveport, LA 71103, USA; Microscopy Core, Research Core Facility, Louisiana State University Health Sciences Center, Shreveport, LA 71103, USA

**Keywords:** HPV16, promyelocytic leukemia nuclear bodies (PML NBs), L1, L2, transport vesicles, primary foreskin keratinocytes, viral DNA, Phospholipase C (PLC γ1, PLC β4)

## Abstract

The interplay between promyelocytic leukemia nuclear bodies (PML NBs) and viral infections is complex and multifaceted, with viruses often engaging in a delicate balance of exploitation and evasion of host immune responses. It is well-established that incoming human papillomavirus (HPV) virions associate with PML NBs; however, the mechanism by which HPV targets PML NBs to facilitate early transcription and replication remains elusive. We have shown that HPV genomes are delivered to the nucleus inside membrane-bound vesicles during mitosis and that PML proteins are recruited to HPV-harboring nuclear vesicles prior to egress. Herein, we demonstrate that the PML isoform II (PMLII), but not other PML isoforms, is selectively attracted to incoming HPV still residing in transport vesicles. Using individual PML NB isoform and HPV16 minor protein, L2, over-expression experiments in HeLa cells, we conclude that PML II binds to the lipids of the HPV harbouring membrane-bound vesicles rather than L2. This is in line with previous observations that PML II is the only isoform binding to lipid droplets in the nucleus. We observed that the association of PML NBs with the HPV16 viral genome was transient and lost over time. We also found that nuclear Phospholipase C β4 and γ1 colocalize with PML II-L1 complexes in infected cells. Our data support a model in which PML II contributes to the establishment of early HPV16 infection by organizing host lipid regulatory factors at incoming viral transport vesicles.

**Author summary:** Promyelocytic leukemia nuclear bodies (PML NBs) are dynamic subnuclear structures involved in antiviral defense, DNA repair, transcriptional regulation and tumor suppression. Despite their antiviral role, HPV exploits PML NBs after nuclear entry, associating its genome with PML proteins, promoting early transcription and infection by shielding the genome from immune responses and degradation. Here, we provide evidence that PML isoform II is preferentially recruited to incoming viral transport vesicles through its interaction with lipids rather than the L2 protein. The association of certain Phospholipases C with the PML II–L1 complex suggests that PML II interacts with lipid-containing HPV16 transport vesicles and may facilitate their PLC- mediated resolution.

## Introduction

Promyelocytic Leukemia Nuclear Bodies (PML NBs) are dynamic, membrane-less subnuclear structures in mammalian cells, formed by self-assembly of PML proteins and components like Sp100, DAXX, and SUMO (1–4). PML NBs play roles in transcriptional regulation, tumor suppression, DNA repair, epigenetic regulation, apoptosis, and antiviral defense (1, 5, 6). Protein recruitment to PML NBs is partly regulated by SUMOylation (7, 8). Seven PML isoforms, generated by alternative splicing, have been identified. PML isoforms I-VI localize to the nucleus and form PML NBs, whereas PML isoform VII lacks the nuclear localization signal and remains cytoplasmic (1). The unique properties and interactions of different PML isoforms with specific proteins highlight the complexity of their regulation within the cell (2). For example, PML II accumulates in the nucleoplasm in close vicinity to newly synthesized lipid droplets and forms distinct lipid associated PML structures (LAPS) that are devoid of SUMO-1, Sp100, and Daxx in contrast to canonical PML NBs. LAPS facilitate the maturation of nuclear lipid droplets that are essential for lipid synthesis (2–4). PML NBs play a dual role in viral infections, supporting host immunity by degrading viral components and activating antiviral pathways. PML NBs regulate transcription through co-regulators like Sp100 and Daxx, suppressing viral gene expression (5–8). However, many DNA viruses, including herpesviruses, adenoviruses, papillomaviruses and polyomaviruses have developed strategies to evade or disrupt PML NBs, often reorganizing them to facilitate early transcription and replication (9–13). HCMV encodes proteins like IE1 and UL83 to degrade PML and disrupt PML NB formation (14–16), while the HCMV tegument protein pp71 facilitates early transcription and genome localization at PML NBs (17, 18). The HSV-1 early protein ICP0 reduces PML SUMO-1 modification (19) and EBV encodes EBNA-LP, which binds PML and inhibits its recruitment to NBs, promoting efficient viral replication (20, 21). Adenovirus E4 Orf3 and E1A-13S oncoproteins disrupt PML NBs by targeting PML II, enabling early transcription (22–24). These disruptions of PML NBs all seem to create a more favourable environment for viral replication and spread.

Human papillomavirus (HPV) and PML NBs interact in a complex manner, with PML proteins exerting antiviral effects that HPV overcomes through evasion strategies. The minor capsid, L2, protein of BPV-1 and HPV16 associates with PML NBs during exogenous expression (25) and in the differentiated layers of natural lesions (26). PML proteins are crucial for early transcription following the delivery of BPV1 genomes and HPV16 pseudogenomes (25) and for establishing HPV16 infection. In PML knockdown HaCaT cells, early transcription is impaired, and the HPV genome is lost after infection (27). These findings highlight the role of PML proteins in shielding the incoming viral genome from innate immune recognition or degradation. Overexpression of the HPV33 L2 protein shows that L2 targets PML NBs and displaces the transcriptional suppressor Sp100 without affecting PML proteins, suggesting a reorganization of PML NBs by L2 to promote early viral transcription (26, 28).

The infectious entry pathway of HPV virions includes preferential binding to extracellular matrix depositions (29–32); conformational changes to both capsid proteins facilitated by host cell chaperones followed by proteolytic cleavage (33–35); insertion of the L2 protein into the endosomal membrane (36–38), and retrograde transport to the trans Golgi network (TGN) mediated by L2 protein (38–40), where the virus resides until the onset of mitosis (41, 42). Finally, HPV is transported along spindle microtubules is essential for transfer to mitotic chromosomes (43). Our laboratory has demonstrated that the HPV genome is delivered to the nucleus of host cells in a membrane-bound vesicle during mitosis (42). This finding was further supported with electron microscopy data provided by Day et al. (44, 45). We further demonstrated that some L1 protein resides within transport vesicles during trafficking and nuclear delivery and is lost when the viral genome is released (46). Recently, we discovered that the viral egress from the transport vesicle is mediated by specific phospholipases C, resolving the lipid component of the vesicle (especially PLCβ4 and PLCδ4) (47). Prior to release, PML protein is recruited to HPV-harboring vesicles through its interaction with viral capsid protein L2 via its SIM motif (48). Notably, PLCγ1, one of the phospholipases C contributing to the phosphoinositide metabolism and nuclear signaling has been reported to associate with PML nuclear bodies, providing a potential link between PML structures and phosphoinositide metabolism (49, 50). In addition, PLCβ4 and PLCγ1 undergo SUMOylation, which can regulate their nuclear localization, trafficking, and signal transduction (51, 52). Despite advance in our understanding of the role of PML NBs in the HPV life cycle, it remains unclear whether any individual PML isoform contributes to HPV infection, what triggers the association of PML NBs with incoming HPV genome transport vesicles and how long this association persists. To address these gaps, we investigated the role of individual PML isoforms in HPV16 infection using knockdown and exogenous expression approaches.

Herein, we report that PML II is preferentially recruited to incoming HPV16 transport vesicles through lipid-associated mechanisms rather than by L2-mediated interaction. PML association with the incoming viral genome is transient and diminished as the cell cycle progressed. We also show that phospholipases PLCβ4 and PLCγ1 associated with PML II–L1 complexes, suggesting that incoming viral transport vesicles provide a platform for PLC-PML II recruitment. Together, these findings suggest that PML II supports the establishment of early HPV16 infection by organizing host factors at incoming transport vesicles and facilitating a postmitotic release of the viral genome.

## Results

### Exogenous PML isoforms colocalize with endogenous PML NBs in HaCaT cells

Knowing the unique properties and interactions of PML isoforms with specific proteins (9), we wanted to determine whether any of the individual PML isoforms are recruited towards incoming viral genomes at the early stage of HPV infection. We overexpressed the individual PML isoform I through VI tagged with EYFP by recombinant lentiviral transduction in HaCaT cells. Overexpression of individual PML isoforms was confirmed by immunofluorescence of endogenous PML and EYFP (Fig. 1). Analysis of these cells indicated that in all cases, 100% of exogenous EYFP-PML isoforms colocalized with endogenous PML (Fig. 1) as well as with Sp100 in the nucleus (Fig. S1). These data were consistent with previous studies demonstrating colocalization of exogenously expressed PML isoforms with PML NBs (1, 50, 53).

**Fig. 1.**
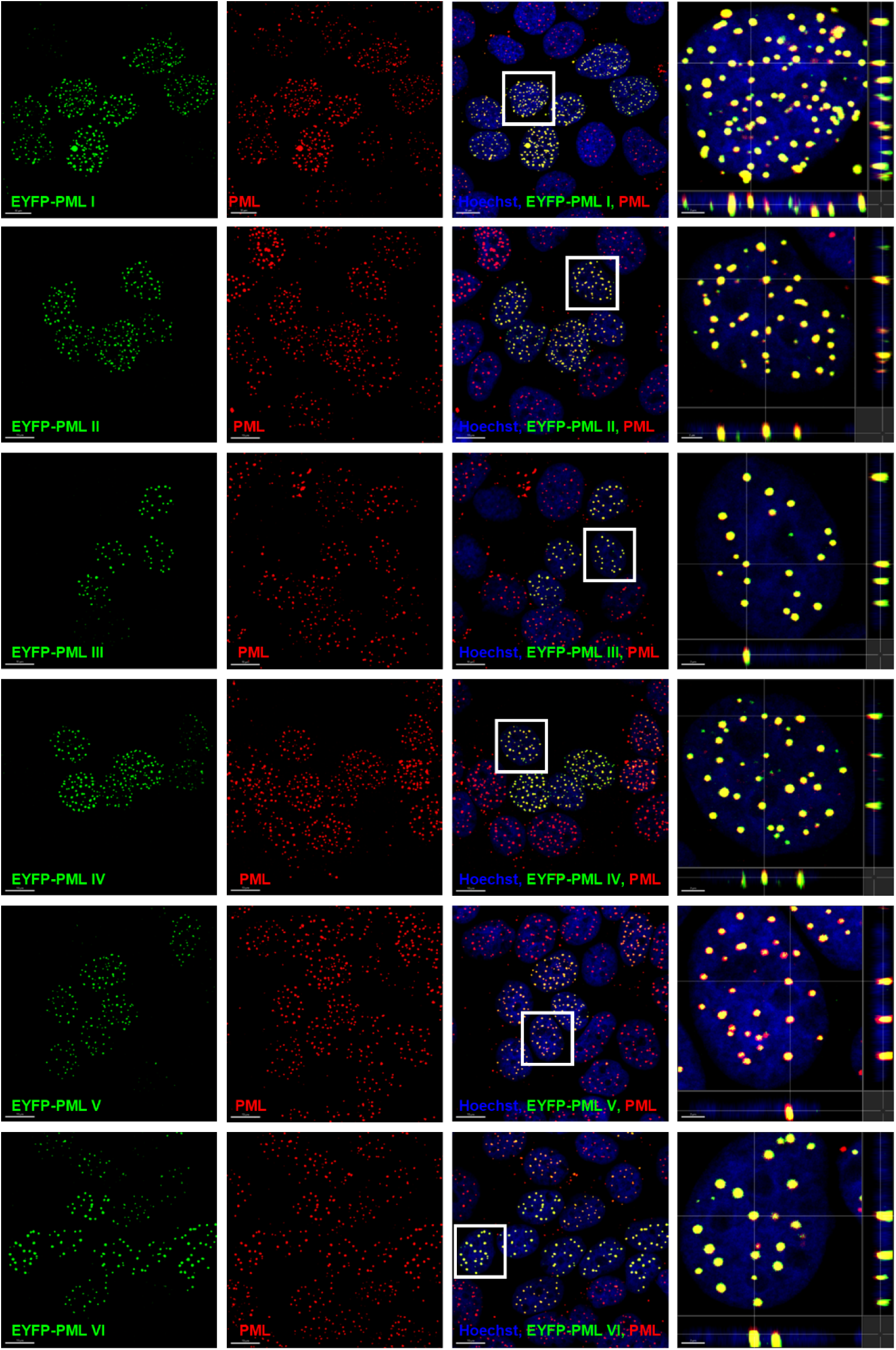
Colocalization of exogenous EYFP-tagged PML isoforms and PML NBs in HaCaT cells. Cells were transduced with lentiviral constructs carrying EYFP-tagged PML isoforms I to VI, selected over six days with G418 (0.5 mg/mL) and stained for PML protein. Confocal images of EYFP-tagged PML (green), total PML (red) and Hoechst 33342 dye for nuclear visualization (blue) are shown as slice. Scale bar 10 µm. Colocalization of overexpressed EYFP-tagged PML isoforms with endogenous PML are visualized as yellow puncta in the merged images, with representative signals displayed in orthogonal cross-section close-up views highlighted on the right-hand side. Scale bar: 2 µm.

### PML II is preferentially recruited to incoming viral transport vesicles

It has been suggested that the PML II isoform accumulates in the nucleoplasm in vicinity to newly synthesized lipid droplets, forming distinct lipid-associated PML bodies (2, 3). We have demonstrated that HPV genome is delivered to the nucleus of host cells in a membrane-bound vesicle during mitosis (42) and that L1 protein resides within the transport vesicle during intranuclear trafficking and is lost upon the release of the viral genome from the vesicle in the nucleus (46, 48). To assess whether PML II has a specific role in the early stages of the HPV lifecycle, we infected HaCaT cells over-expressing individual EYFP-PML isoforms, with HPV16 pseudoviruses (PsVs) for 18 h and stained for L1 protein (Fig. 2A). Quantification of the colocalization between EYFP-PML and L1 showed a significantly increased association of PML II with L1 protein (Fig. 2B). These data demonstrate that PML II is preferentially recruited to incoming viral transport vesicles, distinguishing it from other PML isoforms.

**Fig 2.**
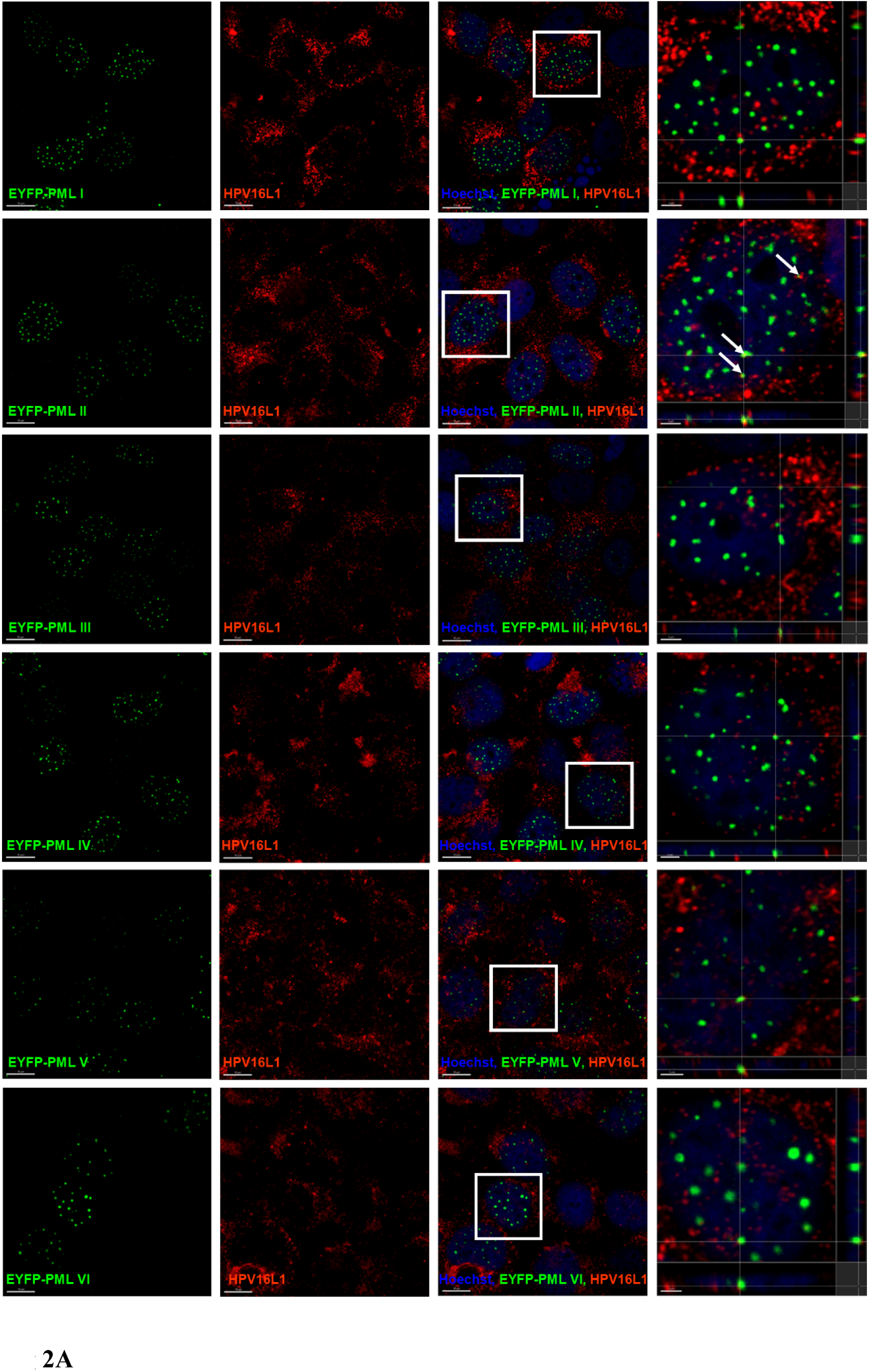
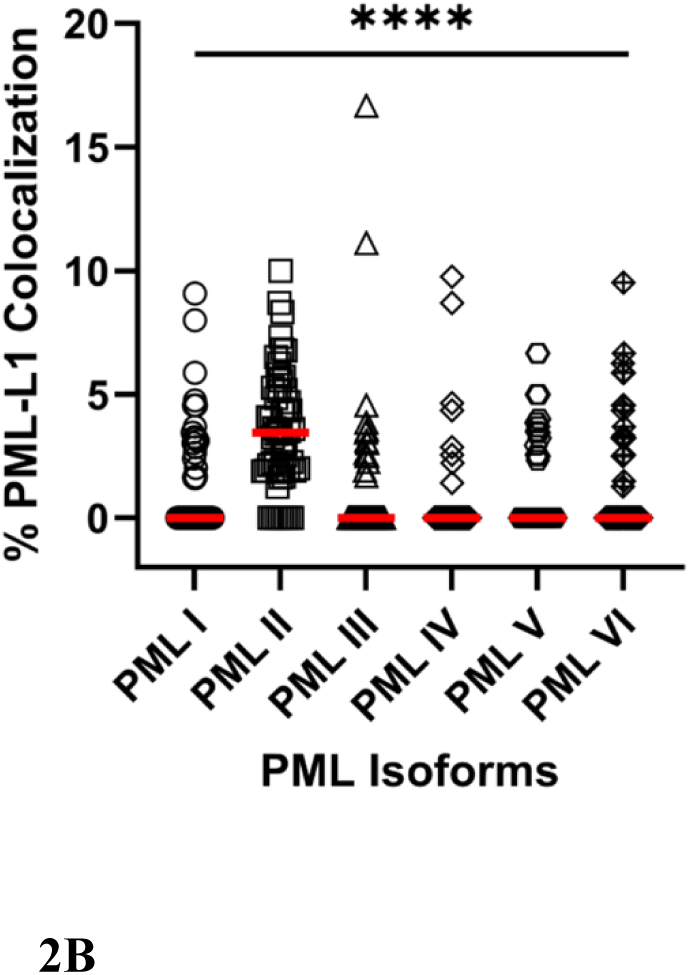
PMLII selectively colocalizes with HPV16 L1 in the transport vesicle in infected HaCaT cells. (A) Cells were stably transduced with lentiviral constructs carrying EYFP-tagged PML isoforms I to VI and selected over six days with G418 (0.5 mg/mL). They were infected with PsVs16 for 18 h and subsequently stained for L1-specific mouse monoclonal antibody (red) and Hoechst 33342 dye for nuclear visualization (blue). EYFP expression (green) was used for detection of exogenous PML isoforms. Yellow puncta (white arrows) in the merged image (right hand side, transverse projection) show the colocalization of overexpressed EYFP-tagged PML II isoforms with L1 protein indicating the presence of the intact transport vesicle. Scale bar: 10 µm. Confocal images are presented as slice. Scale bar: 2 µm. (B) Quantification of EYFP-tagged PML isoforms colocalizing with HPV16 L1. PsVs16 infected HaCaT cell were treated as described previously. The graph shows the percentage of PML isoforms puncta colocalized with the L1 protein per cell. Red line represents median of 40 individual cells analysed per condition from three independent experiments (*N*=3). The P value was determined using the Kruskal-Wallis test. P**** <0.0001.

### Association of PML NBs with HPV16-L2 is independent of specific PML isoforms

Previous studies showed that the L2 capsid protein of BPV-1 and HPV16 associates with PML NBs both after exogenous expression (54) and in differentiated layers of natural lesions (26). Overexpression of HPV33 L2 protein specifically targets PML NBs and displaces the transcriptional suppressor Sp100 from these bodies, without impacting the PML proteins (26, 28). We have shown that the L2 protein remained with the viral genome and associated with PML proteins, transport vesicle resolution and EdU-accessible staining (39, 42, 48). We next asked whether the L2 protein is facilitating this association. We knocked down PML in HeLa cells and confirmed the knockdown by immunofluorescence and Western blot analysis (Fig. 3A-B). PML depleted HeLa cells exhibited a slower growth rate compared to parental cells and prolonged (72 h) lack of PML had detrimental effects on cell viability. For this reason, repopulation of exogenous PML(s) was performed after achieving sufficient knockdown (36 h post-transduction, Fig. 3A-B), and then, cells were selected with puromycin and G418 for 72 h. Western blot analysis showed that all EYFP-PML isoforms displayed multiple bands of unmodified and SUMOylated proteins (Fig. 3C). The expression of reintroduced PML proteins varied across isoforms, with PML I and VI being the most abundant, while PML III and V were the least expressed. PML-isoform repopulated HeLa cells were then transfected with HPV16-L2 expression vector and the localization of the L2 protein with respect to individual EYFP-PML isoforms was assessed by confocal microscopy. We observed that overall, all PML isoforms colocalized with L2 protein to the same extent, 83-90% co-association (Fig. 4A and 4B). These data demonstrated that association of L2 with PML NBs is not dependent on specific PML isoforms and confirmed that recruitment of PML II is not mediated via its interaction L2 protein.

**Fig. 3.**
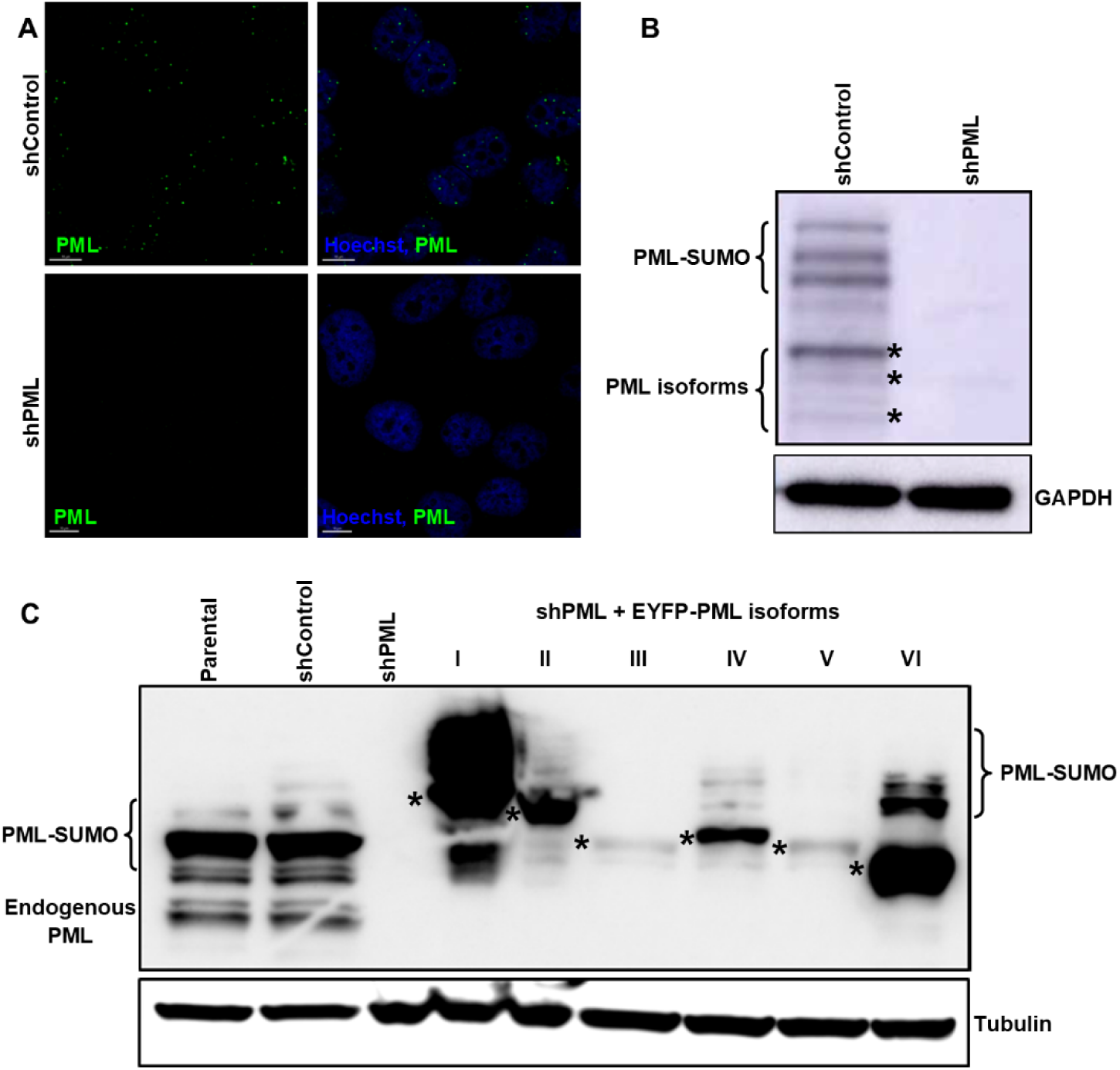
Knockdown of endogenous and overexpression of exogenous PML isoforms in HeLa cells. **(A)** HeLa cells, stably transduced with lentiviruses expressing either scrambled control or PML-targeting shRNA (shPML), were subjected to immunofluorescent staining. Cells were cultured on glass coverslips and stained using an anti-PML antibody (green) and Hoechst (blue) for nuclear visualization. Scale Bars 10 µm. **(B)** Western blot analysis was performed to assess PML protein levels in HeLa cells transduced with either scrambled control or shRNA against PML. Cell lysates were probed with a PML-specific antibody and GAPDH was used as a loading control. The lower bands in the control lane represent the major unmodified forms (marked with asterisks) and their SUMO-modified forms are identified above them. **(C)** Western blot analysis of shPML+EYFP-PML HeLa cell series detected with an anti-PML antibody that detects all endogenous and reintroduced EYFP-tagged PML proteins. The lower bands are major unmodified forms of the PML isoforms indicated with asterisks and their SUMO-modified forms are identified above. Tubulin was used as a loading control.

**Fig. 4.**
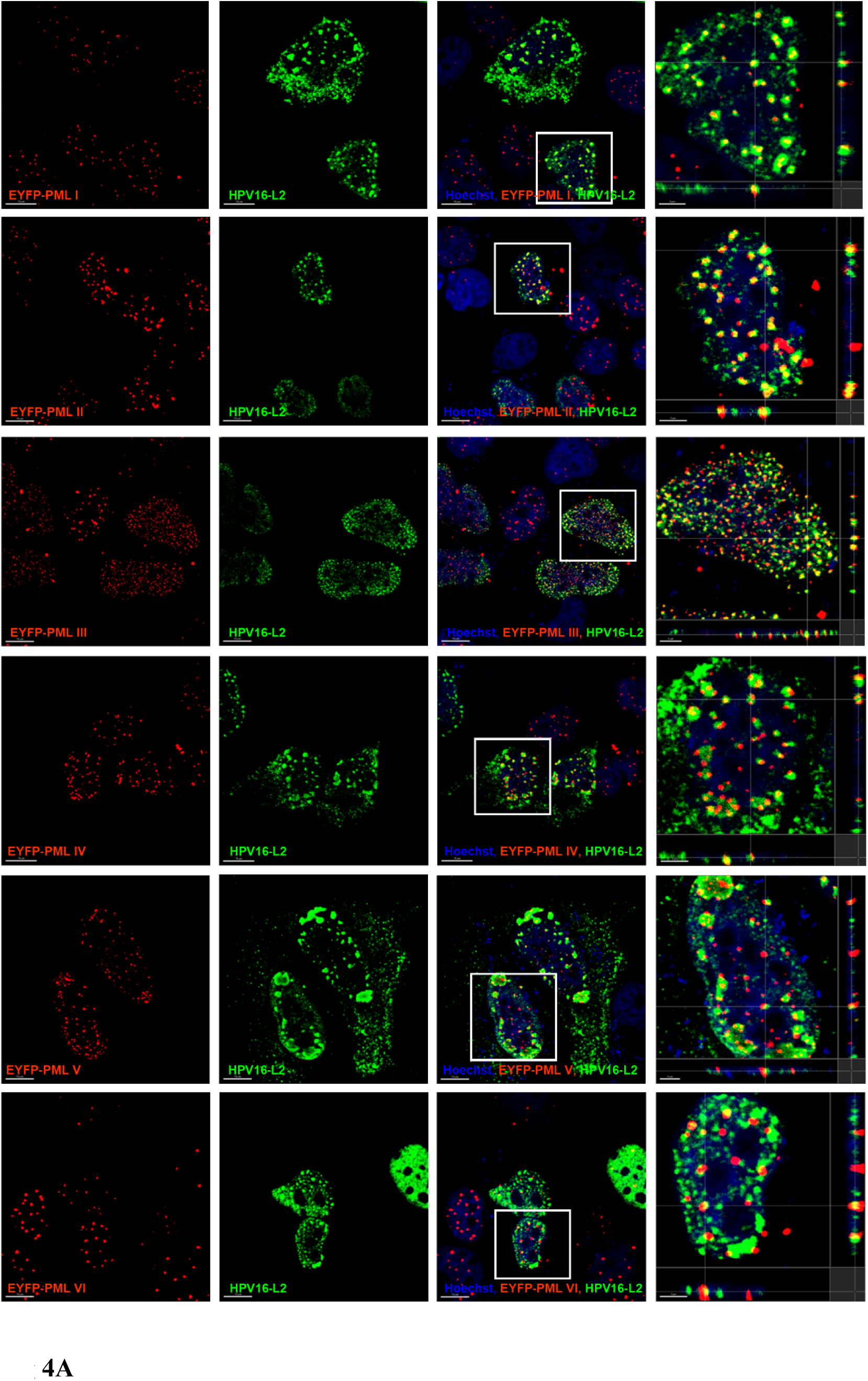
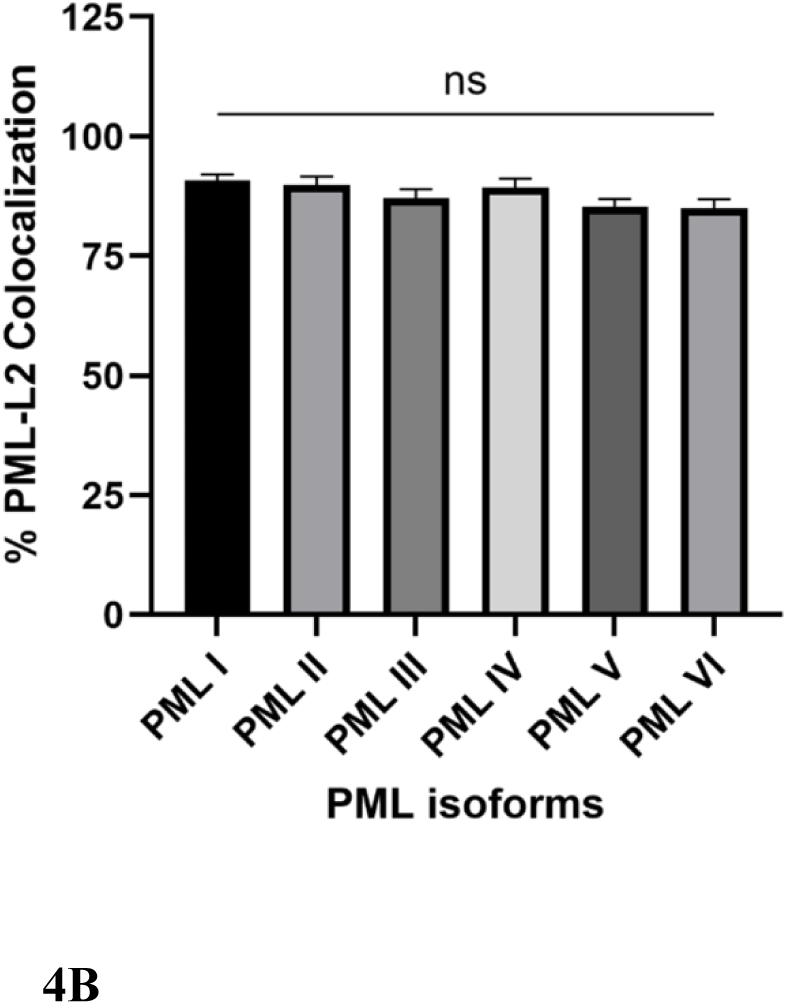
PML NBs association with HPV16 L2 in HeLa cells is independent of specific PML isoform. (A) PML depleted HeLa cells (shPML-HeLa) were transduced with lentiviral constructs carrying EYFP-tagged PML isoforms I to VI and selected over six days with puromycin (2.5 µg/mL) and (G418 (0.5 mg/mL). Cells were cultured on glass coverslips and transfected with HPV16-L2 plasmid for 16 h. Cells expressing reintroduced EYFP-tagged PML isoforms (red) were subsequently stained with L2-specific mouse monoclonal antibody 33L2-1 (green) and Hoechst dye for nuclear visualization (blue). Confocal images are presented as slice. Yellow puncta in the merged image indicates the colocalization of reintroduced EYFP-tagged PML isoforms with L2 protein (Scale bar: 10 µm), with detailed representative signals shown in orthogonal cross-section and magnified view. Scale bar: 2 µm. Images are representative of two independent experiments, 20 nuclei analysed per experiment; *N*=2. (B) Quantification of the average percentage of PML isoforms puncta colocalized with the L2 protein relative to the total PML puncta per cell (n=20 nuclei per each condition from two independent experiments). P value was determined using the One-way ANOVA test. All error bars represent SEM. ns: not significant. Quantifications are representative of two independent experiments, 20 nuclei analysed per experiment; *N*=2.

### Transient association of PML NBs with incoming HPV genomes

We already demonstrated that L1 resides in membrane-bound vesicles and remains associated with the viral genome as PML proteins are recruited in early interphase but is lost in later stages of interphase (48). Based on the observation that PML II associates with lipids and forms lipid-associated PML NBs, we hypothesized that HPV genome association with PML NB is transient and not detectable after the resolution of the transport vesicle and the next mitosis. To test this hypothesis, we used our recently developed infection model where we infect primary human keratinocytes (HFKs) with HPV16 quasivirions (55). HFKs were infected using ECM-to-cell transfer, incubated at several time points (16-72h), and then prepared for HPV16-specific *in situ* hybridization (DNAscope) followed by PML staining. The association of viral genome with PML NBs was analyzed by confocal microscopy. Immortalized HFK cell lines harboring episomal HPV16 genomes were used as a control. At 16 hpi, we observed over 30% of incoming viral DNA (vDNA) closely associated with PML foci. This reflects the interaction between newly delivered HPV-harboring transport vesicles in the nucleus and PML proteins providing favorable conditions for establishment of infection. After one round of mitosis (24 hpi), vDNA-PML association decreases by more than a half (15.3%) and eventually reaches values close to zero at 72 hpi, when cells have undergone approximately three rounds of divisions (Fig 5A and 5B). During infection, most virions are bound to the ECM and are subsequently taken up by cells in an asynchronous manner. Also, cell divisions are not synchronized, giving mixed population of cells in different phases of the cell cycle. This may explain some rare examples of vDNA-PML association at later times postinfection. Therefore, our results suggest that the interaction between viral genome and PML is transient and lost in the subsequent cell cycle, further emphasizing the role of PML in the early stages of viral infection.

**Fig 5.**
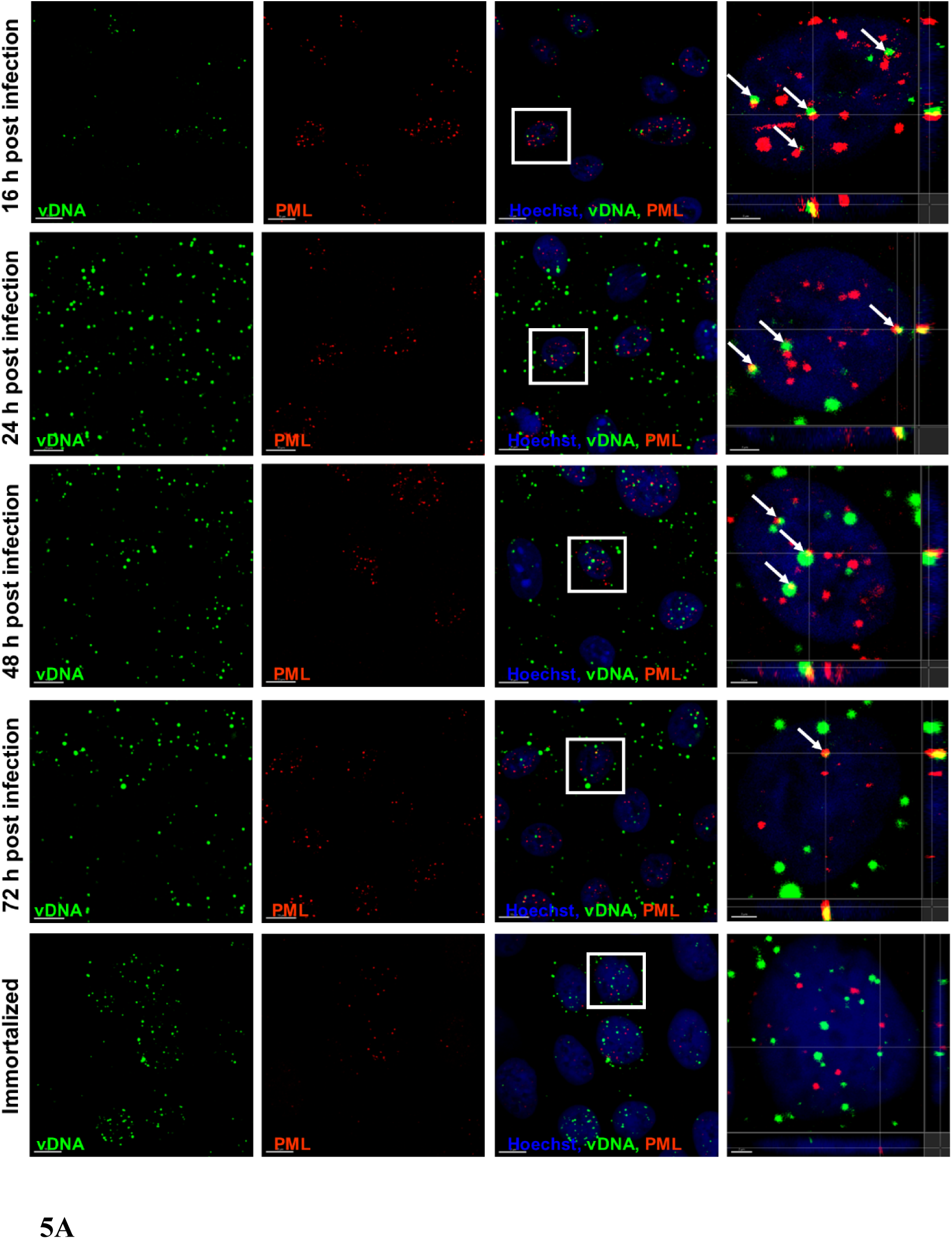
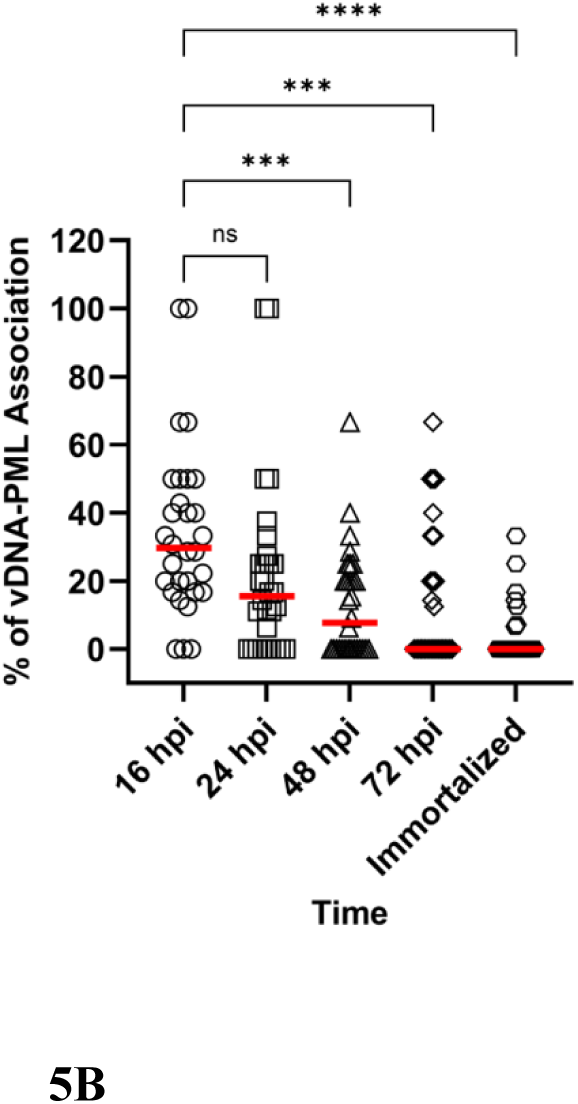
PML NB association with HPV genomes after nuclear delivery is lost over time. (A) Representative images of infected HFKs at 16 h, 24 h, 48 h, 72 h and in immortalized cells. Viral DNA (vDNA) puncta were detected by a DNAscope (green) and cells were also stained for PML proteins (red) and Hoechst dye for nuclear visualization (blue). Confocal images are presented as slices. Yellow puncta with the arrow in the merged image indicate the close association of vDNA with PML protein. Scale bar: 10 µm. Orthogonal cross-section and magnified view on the right shows an association of vDNA and PML in nuclei of infected keratinocytes. Scale bar: 2 µm. Images are representative of 30 nuclei analysed from 3 independent experiments (*N*=3). B) Quantification of the association between PML NBs and vDNA in infected primary foreskin keratinocytes over time. Quantification is based on the percentage of viral DNA puncta detected by DNAscope associated with maximum fluorescence intensity of PML puncta in nuclei of infected HFK. Only cells with nuclear DNA scope signals were included in the analysis. Lines on the graph represent the median (red line) and statistical significance was assessed by the Kruskal-Wallis test for 30 individual nuclei from three independent experiments (*N*=3). P**=0.0005, P****<0.0001, ns= not significant.

### Phospholipases C association with HPV16 transport vesicles and PML II complex

Our recent work has demonstrated that nuclear phospholipases C (PLCs) contribute to HPV16 genome egress from membrane-bound vesicles and proposed that PLCs are recruited to genome-containing vesicles indirectly through L2 or an unidentified mechanism (43). Notably, since PLCγ1 has been reported to associate with PML nuclear bodies, providing a potential link between PML structures and phosphoinositide metabolism (49), we hypothesized that PML II recruits specific PLCs to incoming HPV16 transport vesicles for genome egress. To test this, we depleted endogenous PML in HaCaT cells, repopulated them with EYEP-tagged PML isoforms, as we did previously with HeLa cells, and infected them with HPV16 PsV16s for 24h. These cells were then co-stained for L1 and either PLC isoform β4 or γ1. PLC isoform β4 or γ1 isoform association with PML–L1 complexes was quantified by confocal microscopy. Both tested PLCs isoforms showed significantly greater association with PML II–L1 than with PML I–L1 complexes, the γ1 isoform showed prodigiously more colocalization with PMLII-L1 complexes, roughly 5-6 times more, than the β4 isoform with PMLII-L1 (Fig. 6A–D). Consistent with these findings, PLCβ4 was detected in GFP-mediated PML II co-immunoprecipitation (Fig. 6E), supporting an association between PML II and PLCβ4. Together, these results suggest that PML II preferentially recruits specific PLCs to incoming HPV16 transport vesicles, potentially promoting local lipid remodelling and vesicle destabilization to facilitate viral genome egress in the nucleus post mitosis.

**Fig. 6.**
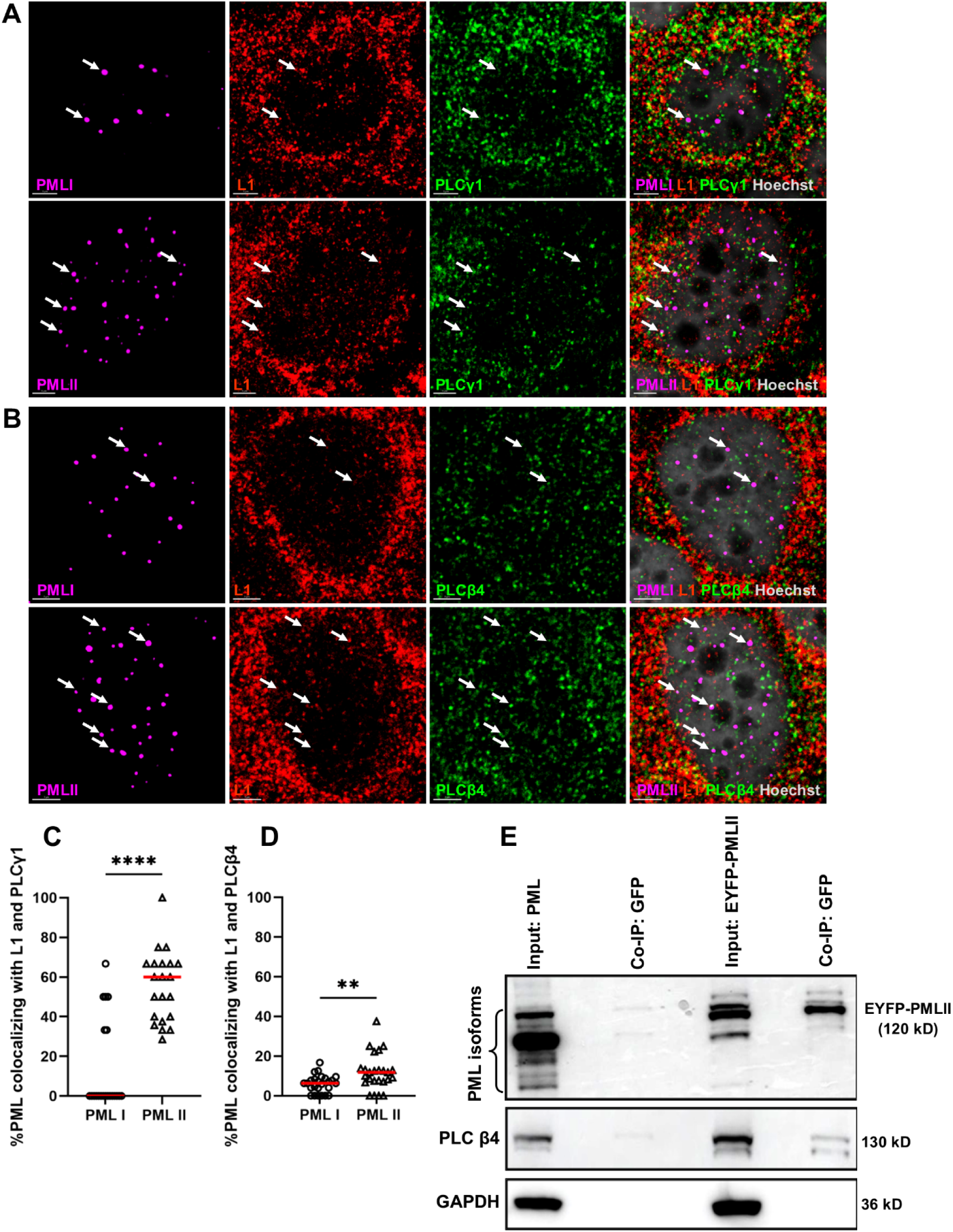
Association of PMLII with PLCγ1 and PLCβ4 on incoming transport vesicle. Cells were stably transduced with lentiviral constructs carrying EYFP-tagged PML isoforms I, and II, and selected over six days with G418 (0.5 mg/mL). They were infected with PsVs16 for 24 h and subsequently stained with a L1-specific mouse monoclonal antibody (red) an antibody against PLCγ1 (**A**), or PLC β4 (**B**) (green) and Hoechst 33342 dye for nuclear visualization (gray). EYFP expression (magenta) was used to detect the exogenous PML isoforms. White arrows in the merged image (right-hand side) indicate the close proximity of overexpressed EYFP-tagged PML II isoforms with L1 and PLC protein. Scale bar: 3 μm. **(C)** Quantification of EYFP-tagged PML isoforms colocalizing with HPV16 L1 and PLCγ1 or PLCβ4 **(D).** The graphs show the percentage of PML isoforms puncta colocalized with L1 in close proximity to PLCγ1/ PLCβ4 protein per cell. Red line represents median of 20 individual cells analyzed per condition from three independent experiments. The P value was determined using Welch’s t-test. P**** <0.0001. **(E)** HaCaT cells expressing EYFP-tagged PML I or PML II were subjected to co-immunoprecipitation (Co-IP) using anti-GFP magnetic beads. HaCaT cells stably expressing scrambled shRNA were included as a control. PML and PLCβ4 were detected by immunoblotting in whole-cell lysates (Input) and GFP co-immunoprecipitates (Co-IP: GFP) using anti-PML and anti-PLCβ4 antibodies, respectively. GAPDH was used as a loading control for whole-cell lysates.

## Discussion

The mechanisms governing the recruitment of PML proteins and their interaction with viral genomes remain largely undefined. Our previous works demonstrated that the incoming viral genomes are delivered to the nucleus of the target cells in membrane-bound vesicular compartments, reside in vesicles throughout the mitosis (39, 42), and the release from the vesicles is delayed until after mitosis and reformation of the nuclear envelope. Further, we demonstrated that PML proteins accumulate around the incoming HPV genome prior to release from the transport vesicles and Sp100 recruitment to the HPV-associated PML NBs is delayed. We also showed that L1 protein is a marker for HPV transport vesicles. This protein directly interacts with the viral genome/L2 complex and remains luminal during intranuclear trafficking; however, unlike L2 protein, it is lost as the viral genome egresses from the vesicle (42, 46, 48). Our findings expand upon the intricate role of PML proteins in HPV infection. Herein, we focused on identifying the specific PML isoforms involved in these early infection stages. Using EYFP-tagged PML isoforms and confocal microscopy, we demonstrated that the PML II isoform, unlike other isoforms, preferentially associates with HPV genome-containing transport vesicles. Since this association is not dependent on L2 protein, we surmise this interaction is mediated by the lipids comprising the HPV genome transport vesicle. Finally, the transient nature of the vDNA-PML interaction suggests that PML proteins act as a temporal scaffold during the early stage of infection.

Using a lentiviral expression system, we demonstrated that all exogenous EYFP-tagged PML isoforms were expressed at endogenous PML levels, distributed uniformly and colocalizing with endogenous PML (Fig. 1) and Sp100 (Fig. S1), consistent with previous findings (1, 49). In PML-depleted HeLa cells, we observed variations in the expression levels of reintroduced PML isoforms, with isoforms I and VI being the most abundant and isoform V the least expressed (Fig. 3C). These differences likely result from post-transcriptional regulation. Isoforms I and II are known to be the most abundant unmodified endogenous PML forms, with PML I-specific mRNA comprising 40–80% of total PML transcripts under normal conditions (54).

The unique characteristic of PML II isoform in both structure and function has been previously demonstrated (2). PML II isoform maturation is associated with lipid droplets. In the nucleoplasm, PML II proteins accumulate around newly synthesized lipid droplets and form distinct lipid associated PML bodies that are relatively devoid of SUMO-1, Sp100 and DAXX differing from the canonical PML NBs (3, 56). Our findings demonstrate that PML II isoform is preferentially recruited to membrane-derived HPV16 transport vesicles containing the viral capsid protein L1. This colocalization distinguishes PML II from other isoforms and suggests a unique functional role for PML II in HPV infection. It is not immediately clear why this particular isoform is recruited to transport vesicles. One plausible explanation is that the PML II isoform that the interacts with lipids on the transport vesicle and forms HPV associated PML bodies. This lipid affinity of PML II appears critical for its role in stabilizing HPV viral genomes, facilitating genome egress. However, further experimentation is needed to test this assumption and to link transport vesicle composition and vesicle resolution mechanism of the incoming viral genome.

Our current study also explored the role of specific PML isoforms in their association with the L2 minor capsid protein. Notably, the L2 protein is known to possess multiple domains that facilitate interactions with cellular factors critical for trafficking (39, 57, 58). These domains include SUMO conjugation and interaction motifs (59, 60), which are pivotal for PML NB formation and protein recruitment. Our previous work demonstrated that L2 protein might recruit PML proteins through SUMO interactions (48). Our current colocalization analyses demonstrated that L2 associates uniformly with all PML proteins, regardless of the specific isoform (Fig 4A, B). This indicates that L2 does not mediate isoform-specific recruitment but rather targets the whole PML NBs (61). These findings suggest that L2 likely employs a distinct mechanism to facilitate its association with carboxy terminal domain (26, 48, 54, 62). Since L2 has no known transcriptional activity for the incoming viral genome, its primary role is likely to localize the DNA to a site conducive to transcription (25). Interestingly, our findings suggest that L2 does not mediate the lipid-driven recruitment of PML II to vesicles. The unique association of PML II with lipids further validates the concept of membrane-bound vesicles and their significance in nuclear trafficking and viral infection.

Considering that the L1 protein resides in transport vesicles and is eventually lost over time after the release of the viral genome from the vesicle (46, 48), we hypothesize that the association of PML NBs with the genome is lost after the subsequent round of mitosis. Herein, we observed transient colocalization of viral DNA with PML NBs. This association was strongest at 16 hpi and gradually declined over 24–72 hpi (Fig. 5A, B) which is consistent with the findings by Stepp et al. (6, 63). The extended (72 h) decline was probably attributed to asynchronous infection, as most virions initially bind to the extracellular matrix (ECM) and are progressively taken up by cells over time. These findings suggest that PML proteins play a critical role during the early stages of infection by interacting with the viral genome. However, this association is not maintained through subsequent cell cycles, emphasizing a temporally restricted role for PML NBs in stabilizing viral genomes and facilitating their early transcription. However, further experiments are required to validate this assumption and to establish a connection between specific lipid interactions that mediate PML II recruitment to transport vesicles and the impact of disrupting PML II recruitment on HPV early transcription and genome stabilization.

PML II is known to be involved in nuclear lipid droplet synthesis (3). Nuclear lipid droplets are enriched in proteins involved in lipid metabolism, including CTP cytidylyltransferase α (CCTα), which has been reported to colocalize with the C-terminal region of PML II (3, 64). Nuclear Phospholipases C are involved in the regulation of phosphoinositide metabolism, and hydrolyse phosphatidylinositol 4,5-bisphosphate (PIP2) to generate diacylglycerol (DAG) and inositol 1,4,5-trisphosphate (IP3). PML nuclear bodies have also been reported to associate with nuclear PLCs, providing a potential link between PML-associated compartments and lipid signaling (47, 49). Interestingly, the PML-II isoform has been shown to be recruited to nuclear lipid droplets and the inner nuclear membrane upon DAG enrichment in membranes, leading to membrane curvature elastic stress (65), providing a potential mechanistic link between nuclear PLC activity, PML recruitment, and HPV transport vesicle resolution. Consistent with these observations, we found that PLC β4 and PLC γ1 associate with PMLII–L1 unlike PML I–L1 complexes (Fig. 6A-D). Moreover, co-immunoprecipitation of PML II with PLCβ4 supports a direct association between PML II and these PLCs (Fig. 6E). Interestingly, we detected PLCβ4 and PLCγ1 colocalizing with PML II in immunofluorescence staining (data not shown) and Co-IP also in the absence of HPV16 infection (Fig. 6E). This finding suggests that PML II and specific PLCs may form a pre-existing cellular complex that is subsequently recruited to incoming HPV16 transport vesicles. Thus, rather than inducing formation of the PML II–PLC complex, HPV16 may exploit a pre-existing host interaction to recruit PLCs to incoming viral transport vesicles. Such recruitment could facilitate local phosphoinositide remodelling and contribute to vesicle reorganization or resolution during viral genome release.

## Materials and Methods

### Cell lines

HEK 293TT, HeLa, and 3T3 J2 fibroblast cells were cultured in DMEM supplemented with 10% FBS, non-essential amino acids, antibiotics, and L-Glutamax. HaCaT cells were maintained in low-glucose DMEM with 5% FBS, antibiotics, and 2.5 mg/mL plasmocin (InvivoGen). Human neonatal primary foreskin keratinocytes (HFKs), obtained from ATCC (PSC-200-010), were cultured in E medium supplemented with 5% FBS, 5 ng/mL mouse epidermal growth factor (EGF, BD Biosciences; 354010), and 10 mM Rho kinase (ROCK) inhibitor Y-27632, which promotes HFK lifespan. Mitomycin-treated mouse NIH 3T3 J2 fibroblasts were used as feeder cells for HFKs, following established protocols (55, 66).

### Generation of HPV16 quasi-and pseudovirions

The pSheLL16 L1/L2 packaging plasmid and the pfwB plasmid, which expresses enhanced green fluorescent protein (GFP), were generously provided by John Schiller (Bethesda, MD). The pEGFP-N1-HPV16 plasmid and pBSCre plasmid were gifts from Jason Bodily (Shreveport, LA). Quasivirions were produced using 293TT cells, following the optimized protocol by Buck and Thompson (67) with minor modifications (66). In brief, 293TT cells were first co-transfected with the pSheLL16 L1/L2 packaging plasmid and the pEGFP-N1-HPV16 plasmid. Twenty-four hours later, the cells were transfected with the pBSCre plasmid. After an additional 48 hours, the cells were harvested, and viral particles were purified as described previously (67, 68). Pseudovirions harboring GFP were produced in 293TT cells, following the protocol described by Buck et al. (67). Viral genome equivalents (vge) were quantified using qPCR of DNA isolated from encapsidated virions extracted with the NucleoSpin® Blood QuickPure kit (Macherey-Nagel; #740569.250).

### Lentivirus transduction

Lentiviral transduction was used to establish HeLa cells with stable knockdown of PML nuclear body (PML NB) proteins and to generate HaCaT or HeLa cells stably overexpressing EYFP-tagged individual PML isoforms. Lentiviral vector plasmids expressing anti-PML (pLKO-shPML), negative control (pLKO-sh-), and PML isoforms (pLNGY-PML I-VI) were kind gifts from Dr. Roger Everett (MRC University of Glasgow Centre for Virus Research) (72). Lentiviral supernatants were produced by co-transfecting HEK-293TT cells with a lentiviral vector plasmid, pMD2.G, and psPAX2 using Lipofectamine™ 3000 reagent (Invitrogen #L3000015). HeLa cells were transduced with lentiviruses expressing shRNAs targeting PML (shPML) or scrambled control shRNAs (ctrl) for 36 h. Cells were selected with puromycin (initially 10 µg/mL, subsequently reduced to 2.5 µg/mL during further passages) (Calbiochem #540222). HaCaT or PML-deficient HeLa cells were transduced with lentiviruses derived from the pLNGY-PML series to express EYFP–PML I to EYFP–PML VI. Transductions were performed in the presence of 8 µg/mL polybrene (Sigma-Aldrich #TR1003-G), and stable cell lines were selected with G418 (initially 1 mg/mL, reduced to 0.5 mg/mL during subsequent passages) (Sigma-Aldrich #G8168).

### HPV16 L2 DNA Transfection

A series of 8 × 10⁴ shPML+EYFP-PML I to VI HeLa cells, cultured on coverslips, were transfected with 0.5 µg of HA-tagged HPV16 L2 overexpression constructs using the Lipofectamine™ 3000 reagent (Invitrogen; #L3000015) following the manufacturer’s protocol. After transfection, the cells were incubated at 37°C for 16 hours to achieve maximum L2 expression. The cells were then fixed with 4% paraformaldehyde and used for PML and L2 immunofluorescent staining.

### Antibodies and reagents

Antibodies used for the study were as follows: PML rabbit (BETHYL; catalog number #A301-167A), PML mouse (Santa Cruz; #sc966), GFP rabbit (ThermoFisher; #A-11122), PLC β4 rabbit (Novus Biologicals; #NBP2-58912), PLC γ1 rabbit (Cell Signalling; #CST 2822S) L1-specific mouse monoclonal antibody 33L1-7 and L2-specific mouse monoclonal antibody 33L2-1 were described previously (42, 67). mAb 33L1-7 recognizes a conserved linear epitope of L1 (residues 303 to 313). We detected the L1 protein by denaturing viral capsid proteins with a Click-iT reaction cocktail before staining (46). Rabbit polyclonal anti-Sp100 antibody was a kind gift from Dr. Hans Will (Heinrich Pette Institute, Hamburg, Germany) (69). Peroxidase-conjugated AffiniPure goat anti-mouse IgG (Jackson ImmunoResearch Laboratories; #115-035-003), and Peroxidase-conjugated anti-rabbit IgG antibodies purchased from Cell Signaling Technology.

### Immunofluorescence staining and microscopy

Cells grown on cover slips at approximately 50% confluency were infected with HPV16 pseudoviruses. Approximately 5-10×10^5^ viral genome equivalents per cover slip were used. In brief, at the indicated times post-infection, cells were washed with PBS and fixed with 4% paraformaldehyde for 15 min at room temperature, washed, permeabilized with 0.5% Triton X-100 in PBS for 5 min, washed, and blocked with 5% goat serum in PBS for 30 min. Cells were incubated for 1 h with primary antibodies at 37°C in a humid chamber, washed extensively and subsequently incubated with AlexaFluor-tagged secondary antibodies for 30 min at 37°C. After extensive washing with PBS, cells were incubated with Hoechst 33342 trihydrochloride trihydrate

(Invitrogen #H3570) for 10 min at room temperature, washed thrice with PBS and mounted in ProLong Glass Antifade Mountant (Invitrogen; #P36984). All IF images were captured by confocal microscopy with a 63x objective using a Leica TCS SP5 Spectral or Leica Stellaris 8 Confocal Microscope. Analysis of colocalization between PML and viral transport vesicle (L1) or L2 in the nuclei was quantified in z-stacks spanning the whole nucleus, otherwise mentioned. Microscopy analyses were performed using IMARIS 10.1.0 software.

### Western blot

Whole-cell extracts were obtained from cell pellets lysed in 1x Laemmli sample buffer supplemented with 2-mercaptoethanol and 1× phosphatase inhibitor Cocktail 2 and 3 (Sigma; #P5726 and P0044, respectively). Proteins were resolved on SDS-PAGE and transferred to nitrocellulose membrane. Membranes were blocked for one hour in 5% Blotting Grade Blocker (BIO-RAD; #170-6404) in 1× PBST and incubated at 4°C overnight with primary antibodies. The anti-PML primary antibodies were used at a 1:2000 dilution. After primary antibody incubation, membranes were washed 3 × 15 minutes in 1× PBST wash buffer. Membranes were then incubated with horseradish peroxidase–tagged goat anti-mouse and goat anti-rabbit secondary antibodies (1:5000) at room temperature for 1 h, then washed 3 × 15 min in 1× PBST. Signals were detected by enhanced chemiluminescence (Thermo Scientific; #1859701 and 1859698). Equal protein loading was confirmed by probing with GAPDH monoclonal antibody (1:4,000) or tubulin - antibody (1:2000).

### Infection using ECM-to-cell transfer

Infection was performed as was published before (55, 66, 70). For secretion of ECM, HaCaT cells were grown on a cover slip for 24 to 48 h until they were confluent. They were incubated in PBS supplemented with 0.5 mM EDTA to detach cells and 8 mg/mL mitomycin for up to 3 h to prevent outgrowth of any residual HaCaT cells. OptiPrep-purified viral particles (0.6 to 1.2 × 10^8^ vge/slide) diluted in 100 µL E medium were added to the ECM for 2 to 3 h at 37°C. This correlates to 200 to 400 vge/cell at 80% confluence. Next, 6 × 10⁴, 5 × 10⁴, 3 × 10⁴, and 2 × 10⁴ low-passage-number primary foreskin keratinocytes were added in E media containing ROCK inhibitor, for harvest at 16 h, 24 h, 48 h, and 72 hpi respectively. After at least 6 h post-seeding of HFKs, 1-1.5 × 10^3^ mitomycin-treated J2 fibroblast feeder cells were added. At the time of fixation, the infected cells reached around 70 to 80% of confluence. In all experiments involving long-term culture, the ROCK inhibitor was removed 7 days post-infection. Immortalized human keratinocytes containing HPV16 episomes were derived from long-term wild-type HPV16-infected HFK cultured in the absence of ROCK inhibitor (55, 66, 71). The episomal or integrated status of the viral genome was assessed using an exonuclease V-qPCR assay (66, 71–73).

### Detection of HPV16 DNA in HPV16-infected HFKs grown in monolayer

HPV16 DNA detection was performed using the highly sensitive hybridization DNAscope assay developed in our laboratory (66, 70, 71, 74). We utilized a Multiplex Fluorescent v2 kit (catalog no. 323100; ACD) and the probe targeting HPV16 L1 (cat no. 315601; ACD). Infected and control cells cultured on coverslips were fixed with 4% paraformaldehyde (PFA) for 15 min at room temperature at 1, 2, or 7 dpi and subjected to the DNAscope procedure. Confocal images were acquired using a Leica TCS SP5 or Leica Stellaris 8 spectral confocal microscope with a 63× objective. Viral DNA puncta and PML foci within nuclei were quantified using Imaris software 4version 10.1.0.

### Coimmunoprecipitation

HaCaT cells overexpressing EYFP-tagged individual PML isoforms I and, II and as a control, sh-control cells were seeded into 6-well plates. Cells were infected with HPV16 PsVs. 24 hpi, cells were harvested and lysed using 0.5% NP-40 Tris-NaCl (pH 7.0) buffer containing 20mM N-ethylmaleimide (NEM) (VWR; #TS15610-0100). GFP magnetic beads (Cell Signaling; #CST 67090) were blocked with 5% BSA for 1 h at 4°C and then incubated with the lysates. After incubation, beads were washed 5× with 0.5% NP-40 Tris-NaCl buffer, and proteins were eluted from beads by boiling at 95°C for 10 min. Proteins in lysates and bead elutes were resolved on 10% SDS-PAGE gels, transferred to nitrocellulose, and probed for the presence of PML, PLCβ, and GAPDH via Western blot.

### Statistical analysis

Data analysis was performed using GraphPad Prism 10.

## Acknowledgments

This research was funded by the National Institute of Health (NIH), grant numbers R01A1164683, P20GM134974, Center of Excellence for Emerging Viral Threats, LSU Health, Shreveport, LA through the Spring 2023 Postdoctoral Fellowship, and LSU Health Shreveport Ike Muslow Postdoctoral Fellowship 2025. We thank Roger Everett, MRC University of Glasgow Centre for Virus Research, for providing the lentiviral vectors for PML knockdown and overexpression. The confocal images presented in this study were generated with the support of the Microscopy Imaging Core at the Research Core Facility of Louisiana State University Health Shreveport RRID:SCR_024775. Mireya Represa-Perez is a fellow supported by the Doctoral Scholarship Program of the Barrie Foundation and LSU Health Shreveport. Lastly, we would like to acknowledge the late Dr. Martin J. Sapp, as this work would not have been initiated or even finished without his guidance and resources.

## Supplemental Material

**Fig. S1.**
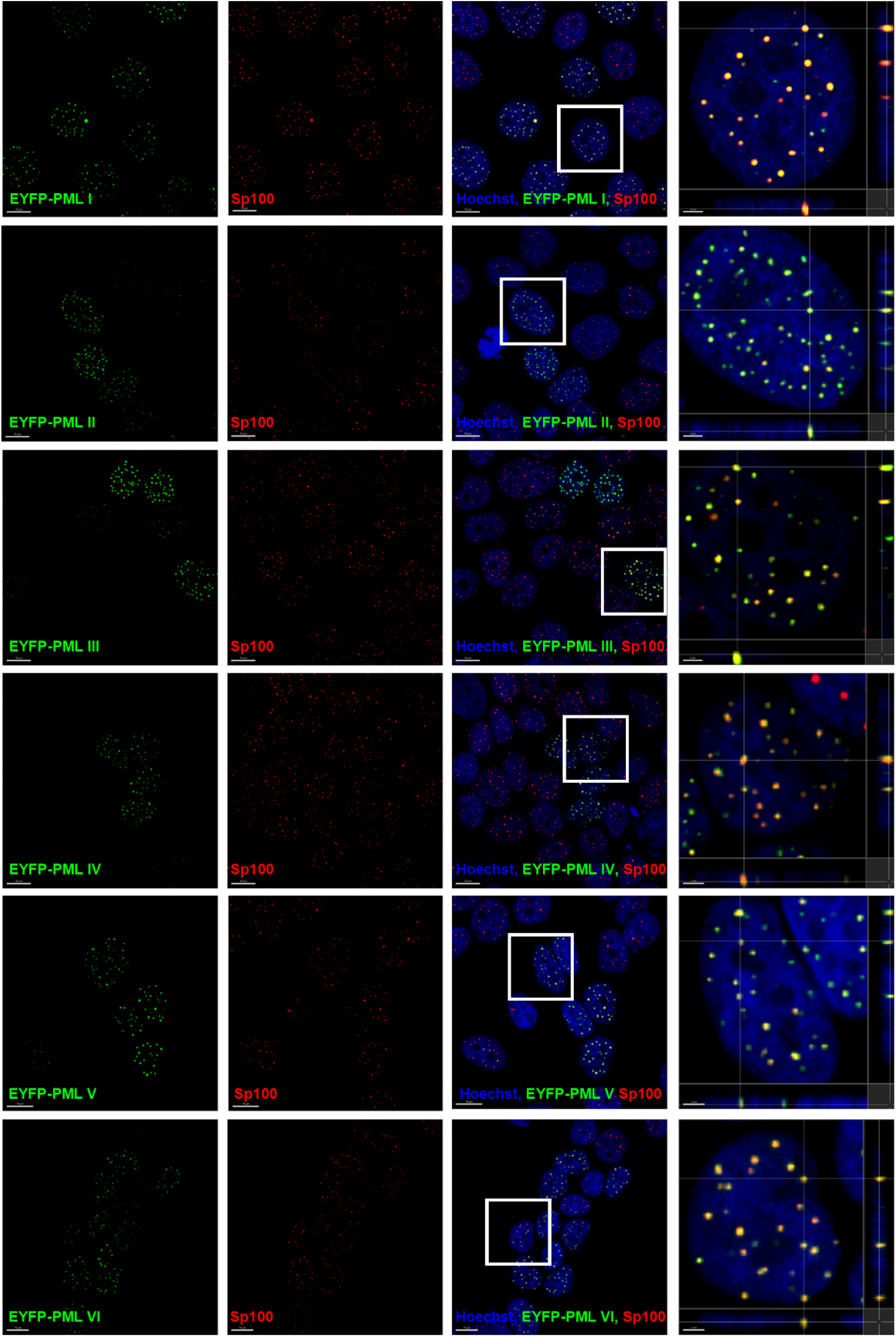
Colocalization of EYFP-tagged PML isoforms and PML NBs component Sp100 in HaCaT cells. Lentiviral constructs carrying EYFP-tagged PML isoforms I to VI were used to transduce HaCaT cells, followed by six days of selection with G418 (0.5 mg/mL). Cells expressed EYFP-tagged PML isoforms (green), stained with anti-Sp100 antibodies to detect the canonical PML NB component Sp100 (red), and Hoechst dye to visualize nuclei (blue). Confocal images are presented as a slice. Colocalization with Sp100 is seen as yellow puncta in merged images Scale bar: 10 µm, with representative orthogonal cross sections and zoomed-in views provided on the right. Scale bar: 2 µm.

